# Meta-analysis of 139 extant *Tara* ocean metagenomes to unveil the relationship between taxonomy and functionality in prokaryotes inhabiting aquatic ecosystems

**DOI:** 10.1101/2020.01.21.913830

**Authors:** Robert Starke

## Abstract

The total microbiome functionality of bacteria was recently predicted to be 35.5 ±0.2 million of KEGG functions. Logically, due to the limitation in space and resource availability of the local community, local functionality will only comprise a small subset of the total functionality but the relationship between taxonomy and functionality is still uncertain. Here, I used a meta-analysis of 139 extant Tara ocean seawater samples from 68 locations across to globe with information on prokaryotic taxonomy on species level from 16S metabarcoding and functionality of prokaryotes on eggNOG gene family level from metagenomes to unveil the relationship between taxonomy and functionality, and to predict the global distribution of functionality. Functional richness showed a statistically significant increase with increasing species richness (P <0.0001, R^2^ =0.64) and increasing species diversity (P <0.0001, R^2^ =0.26) while functional diversity was similar across the different waters, ranging from 2.96 to 3.22. Globally, the highest functional richness was found in the Northern Pacific Ocean and in the North Atlantic Ocean, and decreased at extreme latitudes. Taken together, I unveil the relationship between taxonomy and functionality, and predict the global distribution of functional richness in prokaryotes inhabiting aquatic ecosystems, implying more pronounced effects in terrestrial ecosystems due to larger differences in environmental parameters especially for functional diversity.

---

Ecosystem functioning is mediated by biochemical transformations performed by a community of microbes from every domain of life ^1^. Prokaryotes play key roles in biogeochemical processes such as carbon and nutrient cycling ^2^ and provide the basis for the genetic diversity due to their biomass with 10^4^ to 10^6^ cells per milliliter combined with high turnover rates and environmental complexity ^3^. The visible result of genetic diversity are functions, which can be statistically inferred based upon homology to experimentally characterized genes and proteins in specific organisms to find orthologs in other organisms present in a given microbiome. This so-called ortholog annotation, among others, can be performed in eggNOG ^4^ that comprises 721,801 orthologous groups encompassing a total of 4,396,591 genes and covers all three domains of life (more information about the database can be obtained under http://eggnogdb.embl.de/#/app/home). However, the bottleneck of describing microbiome functions is the low number of fully sequenced and annotated genomes as they are mostly limited to those that have undergone isolation and extensive characterization. Problematically, the vast majority of organisms were not yet studied ^5,6^ and the annotation is based on the similarity to the genomes of the very few studied model organisms. Recently, the total functionality in bacteria were estimated to be 35.5 ±0.2 million functions ^7^ but the relationship between taxonomy and functionality at the local scale and the global distribution of functionality is still uncertain. Here, I used a meta-analysis of 139 extant *Tara* ocean seawater samples using 16S metabarcoding for the taxonomic profile of bacteria combined with metagenome sequencing and eggNOG affiliation for the functional profile of prokaryotes. I aimed to estimate the number of prokaryotic microbiome functions and its Shannon diversity in 20L seawater by identifying the model that best fitted their relationship to species richness and species diversity, and to predict the global distribution of functional richness and functional diversity. I hypothesize that (i) both richness and diversity of local functionality will increase with increasing richness and diversity of prokaryotic species due to the addition of rare functions and (ii) that the functional diversity is similar across different seawater ecosystems as the environments are similar.

In the 139 *Tara* ocean seawater samples enriched in prokaryotes, functionality ranged from 12,328 eggNOG gene families in the Southern Oceans (−61.969° latitude & −49.502° longitude) to 25,238 in the South Pacific Ocean (−8.973° latitude & −139.239° longitude) with an average of 19,523 ±2,682 functions. The relationship between taxonomy and functionality showed statistically significant (P-value <0.05) correlations but the coefficient of determination depended on the specific comparison (Figure 1 & Table 1). The linear relationship of increasing functional richness with increasing taxonomic richness showed the statistically best correlation with a low P-value in combination with a high coefficient of determination (Figure 1a), consistent with my first hypothesis. The addition of new species is likely to add new rare functions ^7^ to the total functional richness which is why an increasing number of species will result in an increasing number of functions. However, this number is limited by space and resource availability of the surrounding environment and its inhabiting microbial community. A maximum of 25,238 functions were carried by 6,254 species but it is likely to assume that seawater samples carry around the average at 19,523 ±2,682 functions in 4,034 ±992 species. Otherwise, the nature of the correlations between taxonomic richness and functional diversity (Figure 1b), taxonomic diversity and functional richness (Figure 1c) and taxonomic diversity and functional diversity (Figure 1d) were all quadratic, implying a local minimum or maximum for each function. Indeed, functional diversity showed a maxima at 3.12 ±0.01 (with 3.11 and 3.13 as 2.5% confidence intervals) with a richness of 5,809 species but a minimum at a functional diversity of 3.08 ±0.01 (3.07-3.09) with a species diversity of 6.4. Functional richness showed a minimum at 17,441 ±446 (16,568-18,317) functions with a species diversity of 6.1. Noteworthy, the increase in functional richness with decreasing species diversity is driven by three samples from the Indian Ocean and their exclusion results in a statistically significant linear and positive relationship (P <0.0001, R^2^ =0.27) which is why I would argue increasing species diversity is increasing functional richness similarly to species richness. Otherwise, functional diversity showed opposing trends for species richness (local maximum) and species diversity (local minimum). Again, the relationship of species diversity is driven by the three samples from the Indian Ocean but also two samples from the Southern Ocean, making it more likely to be a reasonable trend as it was found in different waters across the globe. However, functional diversity ranges only from 2.96 to 3.22 with an average of 3.09 ±0.05 across the 139 seawater samples from different locations where functional richness ranged from 12,328 to 25,238 functions. In comparison, species diversity ranges from 2.48 to 6.97 with an average of 4.03 ±0.99 and a species richness from 2,484 to 6,974. A three-fold larger range in functional richness but a magnitude smaller range in functional diversity suggests, in my opinion, that functional diversity is similar or at least comparable in all the different waters, in line with my second hypothesis. The highest functional diversity reflects both a fit and a healthy community that is able to perform a wide spectrum of possible transformations given by the space and the resource availability of the surrounding environment without overproportioned abundance of singular functions, which would cause a decrease in functional diversity - as seen in the taxonomic data. Environmentally, similar functional diversity across the different waters makes sense as similar processes are performed and the environmental variables such as temperature ^8^, salinity ^9^, oxygen availability ^10^ and dissolved inorganic nutrients ^11^ are similar among the sampled regions. Otherwise, samples from more diverse regions such as the Arctic Ocean or terrestrial ecosystems with a wider range of values of different environmental variables will cause more pronounced differences in functional diversity.

**Table 1:** AICs and pseudo R^2^s of the linear (ln), the logarithmic (lg) and the quadratic (qu) model to describe the relationship between functionality as eggNOG gene families from metagenomes as eggNOG richness and eggNOG diversity to taxonomy as species from 16S metabarcoding and functionality in the form of richness and Shannon diversity. The best fitting model is highlighted in bold.

|  | eggNOG richness vs<br>species richness |  | eggNOG diversity vs<br>species richness |  | eggNOG richness vs<br>species diversity |  | eggNOG richness vs<br>species diversity |  |
| --- | --- | --- | --- | --- | --- | --- | --- | --- |
|  | AIC | pseudo R<sup>2</sup> | AIC | pseudo R<sup>2</sup> | AIC | pseudo R<sup>2</sup> | AIC | pseudo R<sup>2</sup> |
| ln | **2452.71** | **0.64** | -469.09 | 0.14 | 2567.19 | 0.18 | -448.27 | <0.01 |
| lg | 2454.22 | 0.63 | -470.65 | 0.15 | 2568.86 | 0.26 | **-448.19** | **<0.01** |
| qu | 2454.22 | 0.64 | **-470.72** | **0.16** | **2553.80** | **0.17** | -461.90 | 0.11 |

**Figure 1:**
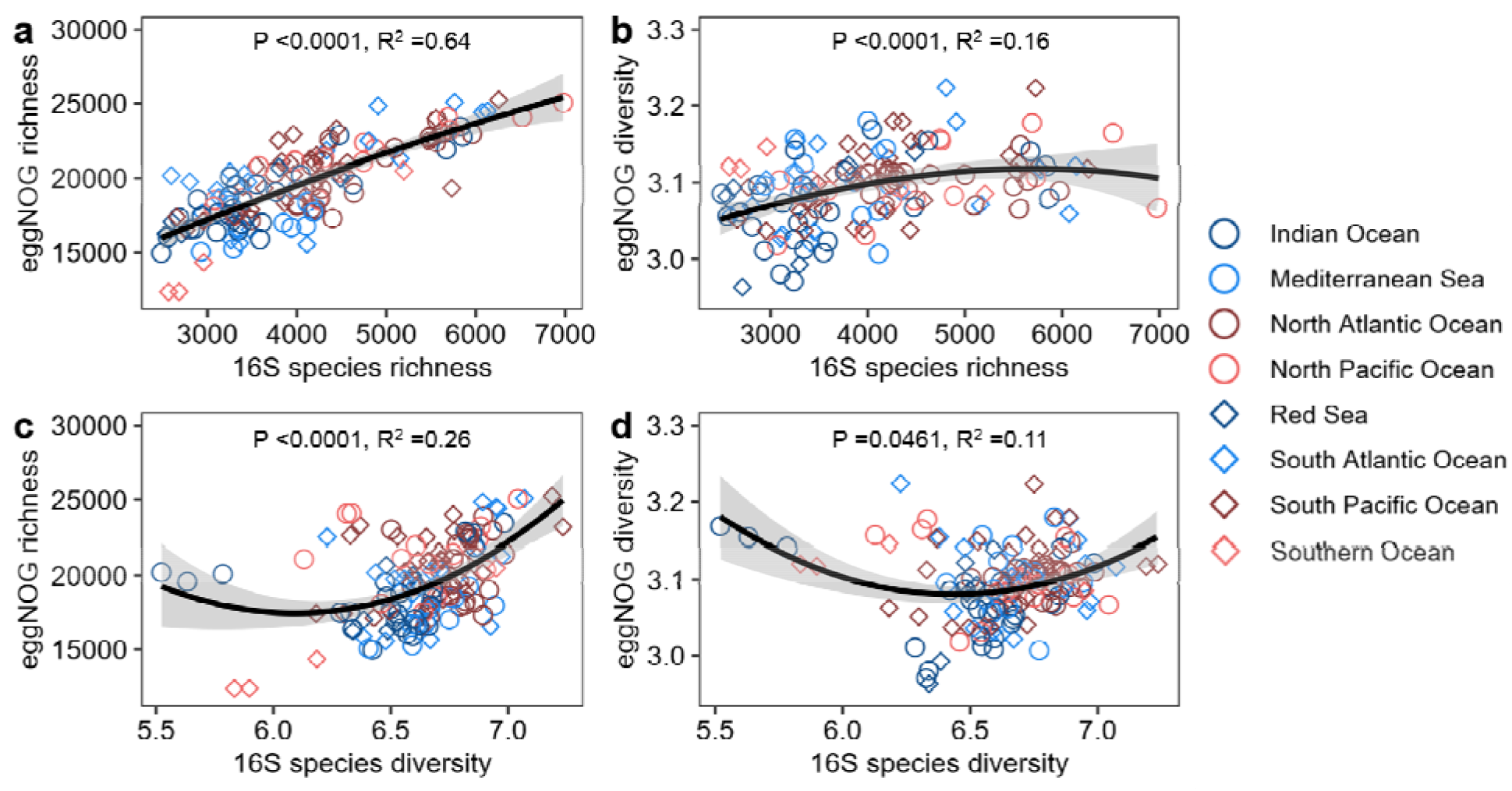
The relationships as smoothed averages between species richness and diversity from 16S metabarcoding and functional richness or diversity of eggNOG functions from metagenomes in 20L seawater samples from 68 locations waters across to globe. The adjusted coefficient of determination (*R*^2^) is given for the best fitting model for each equation: quadratic (a), logarithmic (b), quadratic (c) and linear (d). The P-value was determined by Spearman’s rank correlation.

Globally, functional richness was highest in Northern Pacific Ocean near the American coast and in the North Atlantic Ocean, consistent with statistically significant (P-value <0.05) higher averages of these waters compared to the regions with low functional richness such as the Indian Ocean, the Mediterranean Sea, the Red Sea and the Southern Ocean (Figure 2). Overall, the model comprising second-degree polynomial terms increased in significance (adjusted R^2^ =0.34, P-value =1.105e^−6^) when environmental variables were considered (adjusted R^2^ =0.64, P-value =7.132e^−8^) but nitrate concentration showed the most significant individual effect among the tested environmental variables revealed by the lowest AIC (Table 2) and the highest increase in significance (adjusted R^2^ =0.56, P-value =1.059e^−9^). The correlation between nitrate concentration and functional richness was significant and positively linear (Adjusted R^2^ =0.15, P-value =1.342e^−5^), similar to the significant and positive first-degree polynomial contribution to the best fitting model (P-value =0.000319) to infer high functional richness with high nitrate concentrations. An increase in functionality with changing conditions from aerobic near the surface and anaerobic with increasing depths aligns well with an increasing number of transformation processes and related enzymes involved in microbial respiration. On the one hand, aerobic breathing comprises only one reaction that oxidizes a carbon source to water and carbon dioxide performed by mono- and dioxygenases. Otherwise, the marine nitrogen cycle includes nitrogen fixation by bacteria, nitrate reduction of ammonia production/reduction by phytoplankton in the euphotic zone, followed by sinking/mixing of ammonia and its nitrification to nitrate in the ‘dark ocean’ from where denitrification to nitrogen or vertical mixing with the euphotic zone takes place ^12^. To my surprise as it is contrary to the positive relationship between nitrate concentration and functional richness, low functional richness was found in extreme negative latitudes even though the nitrate concentrations were reported to be highest in these regions with sea-surface concentrations up to 30 mmol N per m^3^ near Antarctica ^11,13^. However, none of these regions of presumably low functionality have actually been sampled and the lower predictions are likely due to the nature of second-degree polynomial functions that forced the maximum of functionality where the samples were taken and result in a minimum towards the extremes. In favor of the lower functional richness near Antarctica are three samples from the Southern Ocean, which align well with the prediction. Admittedly, despite the high coefficient of determination and the significant P-value of the model, most sampling points do not show a close match to the local prediction of functional richness which is why more samples are necessary for a more precise prediction of functionality, especially in the less sampled regions with low functionality such as the Arctic Ocean.

**Table 1:** AICs of the basic model (Lat^2nd^, Long^2nd^) combined with single environmental variables (DE - depths, GT - generation time, Ni - nitrate concentration, Ox - oxygen concentration, Ph - Phosphate concentration and T - temperature) or altogether (Lat^2nd^, Long^2nd^, DE^2nd^, GT^2nd^, Ni^2nd^, Ox^2nd^, Ph^2nd^, SA^2nd^, T^2nd^) to describe the global distribution of functional richness as number of different eggNOG gene families from 139 seawater metagenomes in 1×1 grid cells. In cells containing multiple samples, the sample with the highest functional richness was used (n =74). The best fitting model for the comparison of individual environmental factors to the combination is shown in bold. Then, each combination of first- and second-degree polynomial terms for latitude, longitude and nitrate concentration was tested and the best fitting model shown in bold used to predict the global distribution of functional richness.

| Model (variable<sup>degree</sup>) | AIC |
| --- | --- |
| $\text{Lat}^{2\text{nd}}$, $\text{Long}^{2\text{nd}}$ | 1360.369 |
| $\text{Lat}^{2\text{nd}}$, $\text{Long}^{2\text{nd}}$, $\text{DE}^{2\text{nd}}$ | 1357.532 |
| $\text{Lat}^{2\text{nd}}$, $\text{Long}^{2\text{nd}}$, $\text{GT}^{2\text{nd}}$ | 1362.422 |
| **$\text{Lat}^{2\text{nd}}$, $\text{Long}^{2\text{nd}}$, $\text{Ni}^{2\text{nd}}$** | **1138.57** |
| $\text{Lat}^{2\text{nd}}$, $\text{Long}^{2\text{nd}}$, $\text{Ox}^{2\text{nd}}$ | 1339.934 |
| $\text{Lat}^{2\text{nd}}$, $\text{Long}^{2\text{nd}}$, $\text{Ph}^{2\text{nd}}$ | 1285.342 |
| $\text{Lat}^{2\text{nd}}$, $\text{Long}^{2\text{nd}}$, $\text{T}^{2\text{nd}}$ | 1336.751 |
| $\text{Lat}^{2\text{nd}}$, $\text{Long}^{2\text{nd}}$, $\text{DE}^{2\text{nd}}$, $\text{GT}^{2\text{nd}}$, $\text{Ni}^{2\text{nd}}$, $\text{Ox}^{2\text{nd}}$, $\text{Ph}^{2\text{nd}}$, $\text{SA}^{2\text{nd}}$, $\text{T}^{2\text{nd}}$ | 1115.652 |
| $\text{Lat}^{1\text{st}}$, $\text{Long}^{1\text{st}}$, $\text{Ni}^{2\text{nd}}$ | 1147.529 |
| $\text{Lat}^{1\text{st}}$, $\text{Long}^{2\text{nd}}$, $\text{Ni}^{1\text{st}}$ | 1147.579 |
| $\text{Lat}^{2\text{nd}}$, $\text{Long}^{1\text{st}}$, $\text{Ni}^{1\text{st}}$ | 1137.43 |
| $\text{Lat}^{1\text{st}}$, $\text{Long}^{2\text{nd}}$, $\text{Ni}^{2\text{nd}}$ | 1149.528 |
| $\text{Lat}^{2\text{nd}}$, $\text{Long}^{1\text{st}}$, $\text{Ni}^{2\text{nd}}$ | 1139.336 |
| **$\text{Lat}^{2\text{nd}}$, $\text{Long}^{2\text{nd}}$, $\text{Ni}^{1\text{st}}$** | **1136.7** |
| $\text{Lat}^{2\text{nd}}$, $\text{Long}^{2\text{nd}}$, $\text{Ni}^{2\text{nd}}$ | 1138.57 |
| $\text{Lat}^{1\text{st}}$, $\text{Long}^{1\text{st}}$, $\text{Ni}^{1\text{st}}$ | 1145.579 |

**Figure 2:**
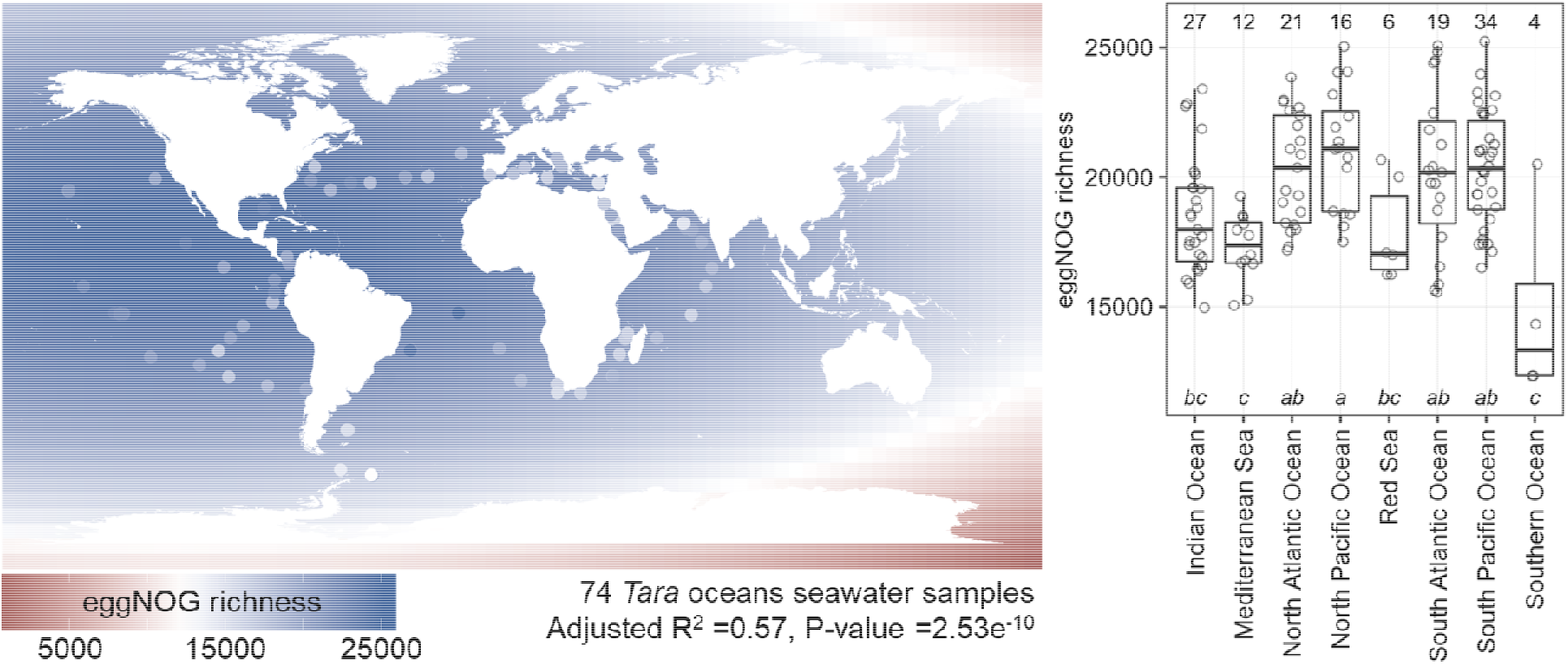
Global distribution of functional richness as eggNOG gene families from metagenomes in 20L seawater samples from 68 locations waters across to globe using non-parametric smoothing for 1×1 grid cells by additive second-degree polynomial models for latitude, longitude, temperature, concentration of nitrate, oxygen and phosphate, and generation time. Including nitrate concentration (AIC =1,138.6) showed the lowest AIC compared to the basic model with latitude and longitude (AIC =1,360.4) and the model with all environmental variables (AIC =1,115.7). The additive pairing of first- or second-degree polynomial terms for latitude, longitude and nitrate concentration showed the lowest AIC value when a first-degree polynomial term is used for nitrate concentration combined with second-degree polynomial terms for latitude and longitude (AIC =1,136.7). The functional richness is also shown based on the region of the different waters with the number of samples as numbers. Groups followed by the same letter are not significantly different according to the HSD test (P-value >0.05).

Altogether, I quantify the relationship between taxonomy and functionality in prokaryotes inhabiting different waters locally and predicted the global distribution of functional richness as functional diversity showed only marginal differences. Noteworthy, the coverage of aquatic ecosystems of the data was admittedly low despite the massive effort of sampling eight oceans over three years but the sampling of more oceans will be beneficial for further predictions. Lastly, due to the grid cell based approach, only half of the bacteria-enriched seawater samples were actually taken into account by the model which is why further expeditions must consider sampling with more spatial separation. Moving forward, this relationship must be examined for terrestrial ecosystems as those generally comprise larger differences in resource availability and environmental variables, potentially resulting in larger differences in functional diversity, as well as for other domains as those govern key roles in terrestrial ecosystem functioning.

## Materials and Methods

### Data collection and correlation between taxonomy and diversity

The publicly available data used to describe the structure and function of the global ocean microbiome ^14^ was downloaded from http://ocean-microbiome.embl.de/companion.html. 139 samples enriched in bacteria comprised the taxonomic profile as annotated 16S OTU count table and the functional profile of prokaryotes as eggNOG gene families annotated to the eggNOG version 3 database ^4^ from the metagenome; both derived from extracted DNA. The richness was determined as the number of different eggNOG gene families or species. The diversity was determined as Shannon diversity *H* according to Equation 1 where *p*_*i*_ is the relative abundance of the eggNOG gene families or prokaryotic species.

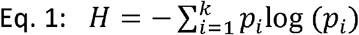

The estimates on functional richness and functional diversity were modelled to species richness as a linear, a logarithmic and a quadratic function using non-linear least squares in the *R* package *nlme* ^15^. The best fitted model was chosen based on the lowest Akaike’s An Information Criterion (AIC) ^16^ with a penalty per parameter set to k equals two. The P-value of their correlation was determined with the function *rcorr* from the R package *Hmisc* ^17^ using the Spearman’s rank correlation. The pseudo coefficient of determination (R^2^) of the non-linear models were estimated with the function *Rsq* in the *R* package *soilphysics* ^18^.

### Global diversity of functional richness

To explore the geographic patterns of functional richness in prokaryotes inhabiting aquatic ecosystems, I assigned the samples to 1×1 degree grid cells covering the globe. Grid-based rather than locality-based analyses can be used to standardize the geographic scale of the analysis, which facilitates cross-region comparisons and limits false presences in the data ^19^. The grid-based approach is broadly favored in biogeographic analyses for its suitability for large-scale comparisons ^20^. In cells containing multiple samples, the sample with the highest number of eggNOG families was selected, resulting in a total number of 74 samples. I used non-parametric smoothing to investigate the changes in functionality (number of eggNOG families) with latitude and longitude in second-degree polynomial terms added to the single or all second-degree polynomial terms of six environmental variables (depth, generation time, nitrate concentration, oxygen concentration, phosphate concentration and temperature). Nitrate concentration combined with latitude and longitude showed the best fit of the data, which was closest to the significance of the model with all environmental variables (Table 2). Then, each combination of first- and second-degree polynomial terms for the three variables was modelled and evaluated. The best fitting model used second-degree polynomial terms for latitude and longitude combined with a first-degree polynomial term for nitrate concentration and was used to predict functional richness in a 5×5 degree grid from −180 to 190 degrees longitude, −90 to 90 degrees latitude and −5 to 45 µmol/L nitrate using the function *dpred* from the R package *iqspr* ^21^. Admittedly, it is questionable that negative nitrate concentration exist but the data was taken as it is available online and since it was present in 28 of 139 samples, their exclusion or further data manipulation could potentially change the data structure. However, it could be the reason for the very low functional richness with extreme latitudes.

## Acknowledgements

I acknowledge my colleague Daniel Morais for showing me the *Tara* ocean data and Petr Capek for his advice on modelling and statistical analysis. This work was supported by the Czech Science Foundation (20-02022Y).

## Contributions

RS designed the study, analyzed the data and wrote the manuscript.

## References

1. Woese, C. R., Kandler, O. & Wheelis, M. L. Towards a natural system of organisms: proposal for the domains Archaea, Bacteria, and Eucarya. Proc. Natl. Acad. Sci. (1990). doi:10.1073/pnas.87.12.4576

2. Falkowski, P. G., Barber, R. T. & Smetacek, V. Biogeochemical controls and feedbacks on ocean primary production. Science (1998). doi:10.1126/science.281.5374.200

3. Whitman, W. B., Coleman, D. C. & Wiebe, W. J. Prokaryotes: the unseen majority. Proc. Natl. Acad. Sci. U. S. A. (1998).

4. Powell, S. et al. eggNOG v3.0: Orthologous groups covering 1133 organisms at 41 different taxonomic ranges. Nucleic Acids Res. (2012). doi:10.1093/nar/gkr1060

5. Pham, V. H. T. & Kim, J. Cultivation of unculturable soil bacteria. Trends in Biotechnology (2012). doi:10.1016/j.tibtech.2012.05.007

6. Martiny, A. C. High proportions of bacteria are culturable across major biomes. ISME J. (2019).

7. Starke, R., Capek, P., Morais, D., Callister, S. J. & Jehmlich, N. The total microbiome functions in bacteria and fungi. J. Proteomics 103623 (2019).

8. Locarnini, R. A. et al. World Ocean Atlas 2013. Vol. 1: Temperature. S. Levitus, Ed.; A. Mishonov, Technical Ed.; NOAA Atlas NESDIS (2013). doi:10.1182/blood-2011-06-357442

9. Zweng, M. M. et al. World Ocean Atlas 2013, Volume 2: Salinity. NOAA Atlas NESDIS (2013). doi:10.1182/blood-2011-06-357442

10. Garcia, H. E. et al. World Ocean Atlas 2013. Volume 3: dissolved oxygen, apparent oxygen utilization, and oxygen saturation. NOAA Atlas NESDIS 75 (2013).

11. Garcia, H. E. et al. World Ocean Atlas 2013, Volume 41: Dissolved Inorganic Nutrients (phosphate, nitrate, silicate). NOAA Atlas NESDIS 76 (2013).

12. Miller, C. Biological Oceanography. (Blackwell Publishing, 2008).

13. Garcia, H. E. et al. World Ocean Atlas 2009, Volume 4: Nutrients (phosphate, nitrate, and silicate). NOAA World Ocean Atlas (2010).

14. Sunagawa, S. et al. Structure and function of the global ocean microbiome. Science (80-.). (2015). doi:10.1126/science.1261359

15. Pinheiro, J., Bates, D., DebRoy, S. & Sarkar, D. nlme: Linear and Nonlinear Mixed Effects Models. R Dev. Core Team (2007). doi: Doi 10.1038/Ncb1288

16. Bertrand, P. V., Sakamoto, Y., Ishiguro, M. & Kitagawa, G. Akaike Information Criterion Statistics. J. R. Stat. Soc. Ser. A (Statistics Soc. (2006). doi:10.2307/2983028

17. Harrell, F. E. & Dupont, C. Package ‘Hmisc’: Harrell Miscellaneous. R Top. Doc. (2016).

18. da Silva, A. R. & de Lima, R. P. Soilphysics: An R package to determine soil preconsolidation pressure. Comput. Geosci. (2015). doi:10.1016/j.cageo.2015.08.008

19. Hurlbert, A. H. & Jetz, W. Species richness, hotspots, and the scale dependence of range maps in ecology and conservation. Proc. Natl. Acad. Sci. U. S. A. (2007). doi:10.1073/pnas.0704469104

20. Větrovský, T. et al. A meta-analysis of global fungal distribution reveals climate-driven patterns. Nat. Commun. (2019).

21. Ikebata, H., Hongo, K., Isomura, T., Maezono, R. & Yoshida, R. Bayesian molecular design with a chemical language model. J. Comput. Aided. Mol. Des. (2017). doi:10.1007/s10822-016-0008-z

